# Host translation machinery is not a barrier to phages that infect both CPR and non-CPR bacteria

**DOI:** 10.1101/2022.11.22.517103

**Authors:** Jett Liu, Alexander L. Jaffe, LinXing Chen, Batbileg Bor, Jillian F. Banfield

## Abstract

Within human microbiomes, Gracilibacteria, Absconditabacteria, and Saccharibacteria, members of Candidate Phyla Radiation (CPR), are increasingly correlated with human oral health and disease. We profiled the diversity of CRISPR-Cas systems in the genomes of these bacteria and sought phages that are capable of infecting them by comparing their spacer inventories to large phage sequence databases. Gracilibacteria and Absconditabacteria recode the typical TGA stop codon to glycine and are infected by phages that share their host’s alternate genetic code. Unexpectedly, however, other predicted phages of Gracilibacteria and Absconditabacteria do not use an alternative genetic code. Some of these phages are predicted to infect both alternatively coded CPR bacteria and standard coded bacteria. These phages rely on other stop codons besides TGA, and thus should be capable of producing viable gene products in either bacterial host type. Interestingly, we predict that phages of Saccharibacteria can replicate in Actinobacteria, which have been shown to act as episymbiotic hosts for Saccharibacteria. Overall, the broad host range of some CPR phages may be advantageous for the production of these phages for microscopic characterization or use as therapy agents, given the current difficulty of CPR cultivation. Absconditabacteria phages and Gracilibacteria phages may have avoided acquisition of in-frame stop codons to increase the diversity of bacteria in which they can replicate.

## INTRODUCTION

Interest in human microbiome-associated Saccharibacteria, Gracilibacteria, and Absconditabacteria (hereafter referred to as SGA) has increased, in part due to their association with disease^1^. SGA are lineages within the Candidate Phyla Radiation (CPR), a monophyletic radiation within domain Bacteria, and are characterized in part by their consistently reduced genomes, small cell sizes, and limited metabolic capabilities^2^. They adhere to lifestyles dependent upon other cells, either by episymbiotic attachment - whereby CPR bacteria attach to, and obtain nutrients from, a larger host bacterium - or by deriving essential compounds such as lipids^3^ from the surrounding microbial community. In most cases, the hosts of CPR bacteria are unknown, but in the case of certain oral and environmental Saccharibacteria, the hosts have been experimentally established to be species of Actinobacteria^4–8^. The attachment by Saccharibacteria can have a profound impact on the Actinobacteria host, leading to cycles of rapid host evolution and drastic changes in host physiology^4,6,9^. The Saccharibacteria-host-bacteria relationship in the human oral cavity has recently been evaluated *in vivo*, demonstrating that in a model of periodontitis, Saccharibacteria reduce the inflammatory effects and pathogenicity of their bacterial hosts^10^. These studies have catalyzed a paradigm shift from the previous characterization of Saccharibacteria as a likely pathogen^11,12^.

In contrast to an episymbiotic lifestyle, one Saccharibacteria species and several Gracilibacteria and Absconditabacteria species are thought to live predatory lifestyles, whereby they feed on specific non-CPR bacteria^6,8,13,14^. Predatory bacteria are an emerging area of research garnering interest as an antibiotic alternative with narrow, targeted effects^15,16^. The predatory Saccharibacteria *Ca. M. amalyticus*, for instance, has been proposed as a tool to precisely consume Mycolata bacteria that are recalcitrant to antibiotic and phage treatments^8^.

Another intriguing feature of Gracilibacteria and Absconditabacteria is that their genomes use an alternative genetic code in which the canonical stop codon, TGA, is instead recognized as glycine (genetic code 25)^17–19^. While the alternative genetic code of Absconditabacteria and Gracilibacteria has been established, very little is known about the genetic code of their phages. It has become clear that phages can adopt a genetic code that is distinct from that of their hosts^18–21^. For example, phages that have reassigned the TAG stop codon to be translated as glutamine infect *Prevotella* that use the standard bacterial code^21^. These alternatively coded phages encode in-frame stop codons within late-stage phage genes to likely prevent premature production of structural and lytic proteins^21,23^. To enable production of these proteins in bacteria that use the standard code, these phage genomes must utilize “code-switching” machinery. These findings raise the possibility that standard coded phages can replicate in bacteria with alternatively coded genomes, but this question has not been investigated to date. Here, we explored the diversity and genomic features, including the genetic codes, of phages that are predicted to replicate in SGA bacteria. In addition to the basic science motivation, phages that replicate in SGA bacteria could have practical importance, as phages can be used to alter the composition of microbiomes with species or strain specificity^24,25^.

## RESULTS

### CRISPR-Cas Systems within Saccharibacteria, Gracilibacteria, and Absconditabacteria

As CRISPR spacers are fragments of phage genomes stored within CRISPR-Cas systems, a common technique used to link phages to their bacterial hosts is via spacer-phage matching^26–29^. To find CRISPR-Cas systems encoded within SGA bacteria, we began with a previously compiled database that contained 862 genomes from the Saccharibacteria, Gracilibacteria, and Absconditabacteria (SGA) lineages^30^ (Supp. Table 1). SGA bacteria in this database are from a wide array of environments, including human microbiome, non-human animal microbiome, soil, freshwater, and marine ecosystems. We de-replicated the database at 99% average nucleotide identity (ANI) to form a non-redundant database of 391 genomes (318 Saccharibacteria genomes, 46 Gracilibacteria genomes, and 27 Absconditabacteria genomes) (Supp. Table 2).

To survey the incidence of complete CRISPR-Cas systems within our genomic database, we searched for Cas loci using the full suite of Tigrfam HMM profiles (*Materials and Methods*) within the genomes that contained high-confidence CRISPR arrays predicted by CRISPRCasFinder (CCF). We manually examined scaffolds that contained Cas gene annotations to ensure that they originated from one of our three SGA lineages (Supp. Table 3). We identified 43 CRISPR-Cas systems present in our non-redundant database (Figure 1A, Supp. Table 4). Encoding at least one CRISPR-Cas system were: 16 Gracilibacteria genomes (among 46 genomes - 34.8% prevalence), 22 Saccharibacteria genomes (among 318 genomes - 7.9% prevalence), and 2 Absconditabacteria genomes (among 27 genomes - 7.4% prevalence). The 34.8% prevalence of CRISPR-Cas systems among Gracilibacteria genomes is substantially higher than reported rates for other CPR^31^ bacteria and closer to the typical CRISPR-Cas system prevalence across the domain Bacteria (∼39%)^32,33^.

**Figure 1.**
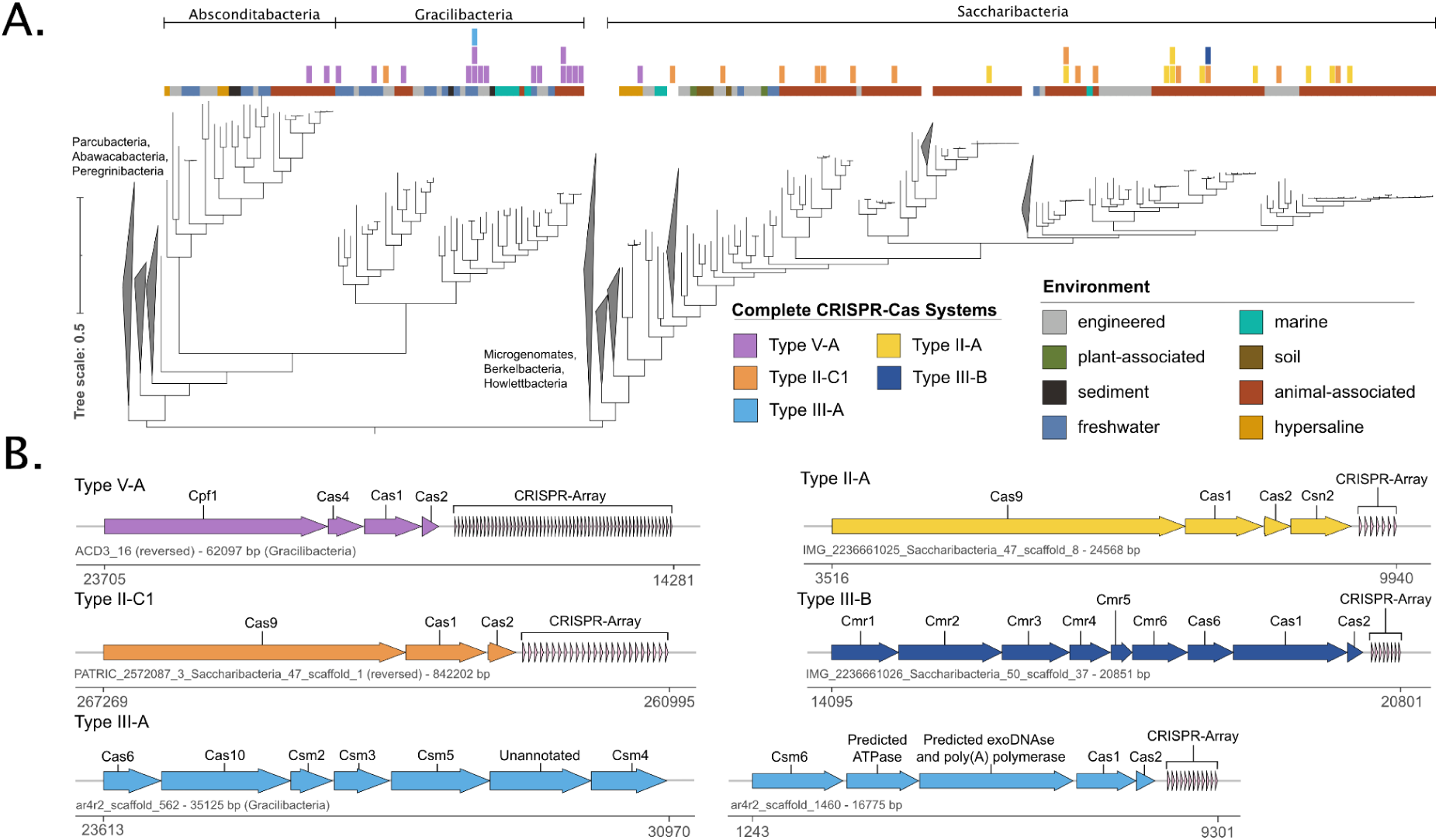
Distribution of CRISPR-Cas Systems in SGA bacteria. **A**. Maximum-likelihood tree based on 16 concatenated ribosomal proteins (*Materials and Methods*). The identified CRISPR-Cas systems and the environmental origin of genomes are overlaid above. **B**. Gene architecture of representative CRISPR-Cas types identified in our SGA database. Color corresponds to the system types displayed in A. Below the gene diagrams are the name and size of the scaffold encoding the system, along with the chromosome coordinates of the system.

When comparing the environmental origin of the genomes containing CRISPR-Cas systems, there is an apparent discrepancy in system distribution between the three SGA lineages. In Saccharibacteria, as has been previously observed^30,34,35^, CRISPR-Cas systems are abundant in animal-associated environments – human and other mammal microbiomes – and scarce in other environments. Only 3 of the 25 CRISPR-Cas systems in Saccharibacteria from our database are from non-animal-associated environments. In contrast, 11 of the 16 Gracilibacteria CRISPR-Cas systems belong to genomes from non-animal-associated environments (Figure 1A).

Despite the streamlined nature of CPR genomes, we also identified five genomes that encode multiple CRISPR-Cas systems. Remarkably, a Gracilibacteria genome (ALUMROCK_MS4_BD1-5_24_33_curated) encodes three distinct Cas loci, including a CRISPR-Cas array with 80 spacers. When Cas genes and CRISPR arrays are taken together, this specific genome dedicates 24,788 bp of its 2,138,004 bp genome (1.16%) to CRISPR-Cas defense systems.

We examined the architecture of our complete CRISPR-Cas systems and categorized the systems, based on previous classifications^32,36^, into five distinct CRISPR-Cas subtypes: type II-A, type II-C1, type III-A, type III-B, and type V-A (Figure 1B). The distribution of the CRISPR-Cas subtypes in relation to SGA lineage is as follows: Saccharibacteria encode type II-A, type II-C1, and type III-B systems; Gracilibacteria encode type V-A and type III-A systems; Absconditabacteria encode type V-A systems. There were two exceptions to these generalizations: 1) one Saccharibacteria genome encodes a type V-A system and 2) one Gracilibacteria genome encodes a type II-C1 system. 4 of the 5 subtypes have been previously identified in CPR bacteria, type II-A^37^, type II-C1^34,37^, type III-A^38^, and type V-A^13^. To our knowledge, subtype III-B has not been previously reported in CPR bacteria. While we expected to find primarily class 2 CRISPR-Cas systems, which are typically more compact and utilize a single effector gene^32,36^, the type-III systems (Class 1) we identified utilize a multisubunit effector complex. The type III-A and type III-B systems we identified, for instance, contained nine identifiable Cas genes together in a single operon. The targets of these subtypes are known to vary: type II and type V-A systems are thought to target double-stranded DNA, while type III-A and type III-B systems are capable of targeting both DNA and RNA^32,36^. This may indicate that some Saccharibacteria are capable of targeting both DNA and RNA phages. Interestingly, we also found that Gracilibacteria and Absconditabacteria almost exclusively rely on type V CRISPR-Cas systems, despite the system’s rarity among bacteria (<2% of all CRISPR-Cas systems identified in bacteria)^36^. Further, we compared the system architecture within each subtype based on average amino acid identity (AAI) of component proteins and noted a mostly uniform architecture within each subtype (Supp. Figs. 1-3). Among the systems, we found 10 variants of the canonical CRISPR-Cas subtypes that contained unannotated ORFs in the interior of a Cas operon. These variants may represent novel subtypes within the broader system classification. One of these variants appears to be a type II-A system (Supp. Figure 1), and nine appear to be type V-A systems (Supp. Figure 3). The functions of these unannotated ORFs and whether they participate in concert with their respective CRISPR-Cas systems remain topics of future study.

To evaluate the novelty of the annotated genes within the complete CRISPR-Cas systems, we compared each Cas sequence to NCBI’s nr database (Supp Figure 4). While most Saccharibacteria and Absconditabacteria Cas genes are well-represented in Genbank, there were a number of our Gracilibacteria Cas genes with less than 50% amino acid identity to known sequences. One such gene is a *cas9* gene that only displays a 34% amino acid identity to the best match in Genbank.

### Putative SGA-infecting Phages

To identify potential phages that infect the SGA bacteria of our database, we extracted 1,296 non-identical spacers from our quality-controlled, high-confidence arrays from the complete genome dataset (147 arrays encoded in 119 scaffolds) (Supp. Table 3). We also searched metagenomic reads for variant sequences that are not reflected in the consensus metagenomic assembly (*Materials and Methods*). We recovered an additional 344 unique spacers from 10 SGA genomes.

To identify putative phages of SGA bacteria, we compared our set of 1,640 spacers to two large phage databases, IMG/VR^26^ and GVD^39^ using the thresholds of at least 95% coverage and less than two mismatches (Supp. Tables 5, 6). After dereplicating hits at 99% ANI, we identified 547 distinct phage scaffolds that putatively infect SGA bacteria (Supp. Table 7). Based on spacer-matching, 440, 57, and 50 of our identified phages were predicted to infect Saccharibacteria, Gracilibacteria, and Absconditabacteria, respectively. Additionally, 26 of the 547 phage genomes were circularized (Figure 2A, Supp. Table 8). We further identified 147 integrated prophages from the same set of de-replicated SGA genomes: 120, 15, and 12 prophages within Saccharibacteria, Absconditabacteria, and Gracilibacteria, respectively (Supp. Table 9).

**Figure 2.**
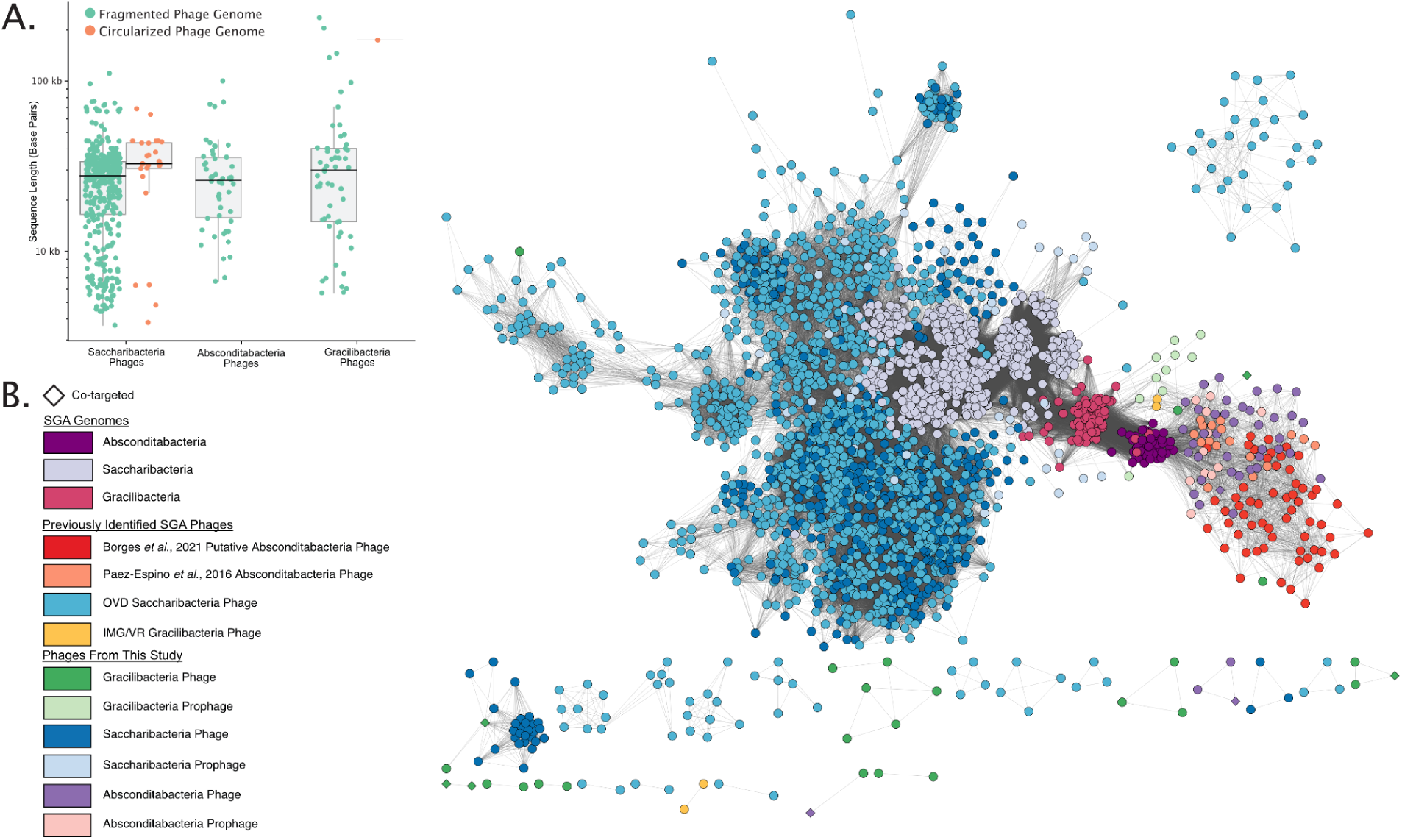
SGA phage genome size and protein-sharing network analysis **A**. Size and completeness of the putative SGA phages. **B**. Protein-sharing network of the putative SGA phages and SGA bacterial genomes, where each node represents a phage or bacterial genome. Nodes are clustered together based on protein similarity and number of shared proteins. Previously identified SGA phages^23,40,41^ were included in the network. Nodes are colored based on the predicted host of the phages or the SGA genome taxonomy. Co-targeted phages indicate those targeted by spacers from CRISPR-Cas arrays of both SGA and non-SGA bacteria.

We characterized our putative SGA-infecting phages and their hosts by generating a protein-sharing network in which the proteomes of phages and SGA hosts are clustered based on similarity (Figure 2B). The proteomes of the Saccharibacteria phages and Absconditabacteria phages predicted in this study cluster with those of their predicted host bacteria and those of previously identified phages predicted to infect their host bacteria, strongly supporting host inference based on our spacer targeting analyses. When clustered in a separate network with non-SGA reference phages, the SGA phages predicted in this study tended to cluster apart from the non-SGA reference phages (Supp. Figure 5). This includes several phages newly predicted to infect Saccharibacteria, which form distinct clusters apart from previously identified Saccharibacteria phages or non-CPR reference phages, and are thus inferred to be novel lineages. Within both networks, our putative Gracilibacteria-infecting phages did not form a central cluster. Within the network between SGA bacteria and their predicted phages, three Gracilibacteria phages predicted by spacer matching cluster with Gracilibacteria prophages or Absconditabacteria phages (Figure 2B). Within the protein-sharing network containing non-SGA references phages, a number of Gracilibacteria phages place within a sparse network that includes reference phages predicted to infect bacteria from either the Bacteroidota or the Firmicutes Phyla (Supp. Figure 5).

### Diverse coding strategies among Absconditabacteria phages and Gracilibacteria phages

To investigate the genetic codes of our identified phages, we predicted ORFs for each phage genome larger than 20 kb in both the alternative code 25 (the genetic code of Gracilibacteria and Absconditabacteria) and the standard code 11. Using these predictions, we calculated the coding density (portion of the genome dedicated to coding genes) in each code. Differences in coding densities between code 25 and code 11 were negligible for Saccharibacteria phages, indicating that they share genetic code 11 with their putative hosts (Figure 3A, Supp. Table 10). Contrary to our expectations, 34 of the 38 putative Gracilibacteria-infecting phages larger than 20 kb displayed small changes in coding density between the two genetic codes, indicating that they are not clearly alternatively coded (Figure 3B, Supp. Table 10). Most putative Absconditabacteria-infecting phages displayed a much higher coding density in code 25 compared to code 11, indicating that they mainly share their host’s alternative genetic code (Figure 3C, Supp. Table 10). However, 6 of the 50 Absconditabacteria phages are not clearly alternatively coded (less than a 10% change between code 25 and code 11 coding densities). Notably, Gracilibacteria prophages and Absconditabacteria prophages displayed a much higher coding density in code 25, indicating they preferentially adhere to the alternative genetic code 25.

**Figure 3.**
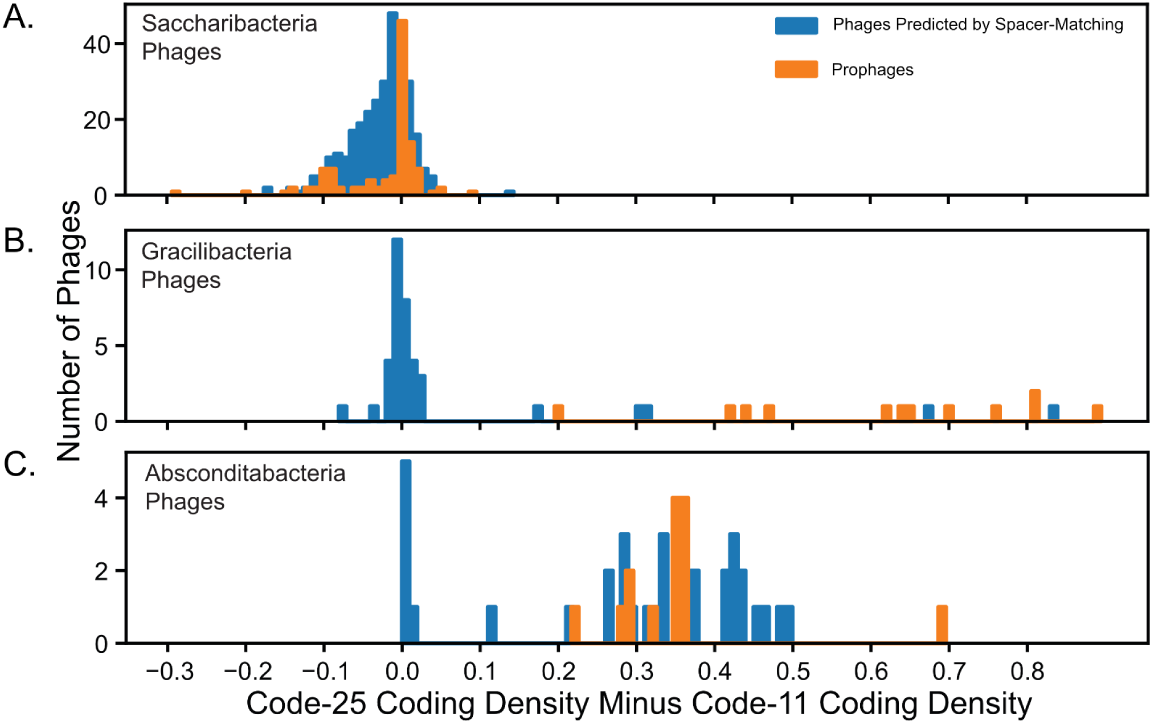
Phage genetic code analysis. Histogram of phages displaying the change in coding density between code-25 and code-11 predictions for phages of (A) Saccharibacteria, (B) Gracilibacteria, and (C) Absconditabacteria. A larger x-value indicates a higher likelihood of adhering to genetic code 25, while an x-value near zero indicates the likely usage of genetic code 11. Only phages larger than 20 kb were included in this analysis.

To further assess the genetic code of the putative Gracilibacteria phages and Absconditabacteria phages, we annotated and visualized the predicted ORFs of each phage in both code 11 and code 25. Examination of 40 Gracilibacteria and Absconditabacteria phage genomes that were not clearly alternatively coded showed that they had very similar gene annotations and genome architectures in both genetic codes (Figure 4A, Supp. Table 11). Further, their genes displayed an absence of in-frame TGA codons and the presence of multiple, different stop codons in close proximity at gene termini (Figure 4A, Supp Table 11). These phage genomes are therefore likely compatible with both code 11 and code 25. They contrast with the clearly alternatively coded Gracilibacteria phages and Absconditabacteria phages, which contained high densities of in-frame TGA codons and displayed almost no gene annotations in code 11 (Figure 4B).

**Figure 4.**
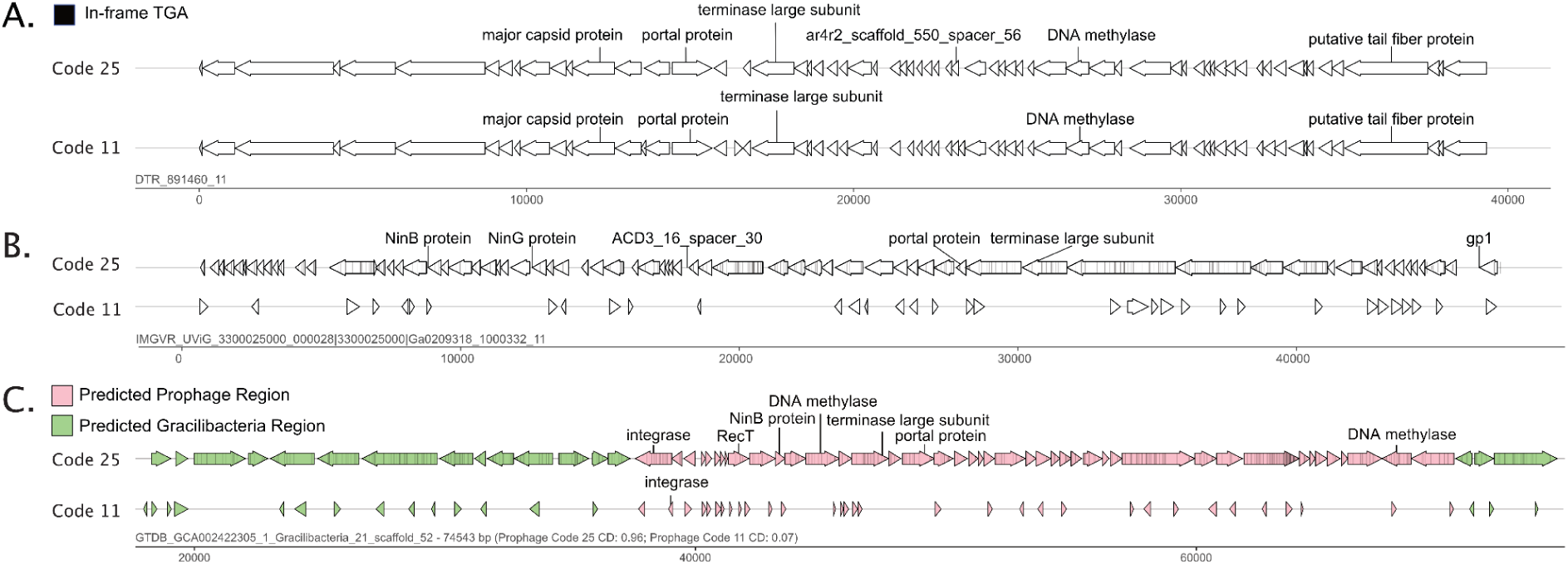
In-frame TGA codon usage among putative Gracilibacteria phages and prophages. **A**. Genome diagrams in code 25 and code 11 of a code 11-compatible Gracilibacteria phage. In-frame TGA codons are marked by a black line. **B**. Genome diagrams in code 25 and code 11 of a clearly alternatively coded (code 25) Gracilibacteria phage. **C**. Genome diagrams in code 25 and code 11 of a Gracilibacteria prophage. Predicated prophage regions and genome regions are colored pink and green, respectively.

Notably, all Gracilibacteria prophages and Absconditabacteria prophages contained ORFs with high densities of in-frame TGA-codons. The identification of these genome regions as prophage rather than novel portions of bacteria genomes is supported by the identification of phage genes that produce a portal protein, tail-related protein, terminase, or integrase (Figure 4C, Supp. Table 12). We conclude that the Gracilibacteria prophages and Absconditabacteria prophages are likely exclusively compatible with code 25.

As it was surprising to find Gracilibacteria phages and Absconditabacteria phages that are compatible with code 11, we sought to further verify that these phages infect their putative alternatively coded hosts. Three Code 11-compatible Gracilibacteria phages and Absconditabacteria phages in the protein-sharing network (Figure 2B) cluster with clearly alternatively coded Gracilibacteria prophages or Absconditabacteria phages (Supp. Figure 7). Additionally, we predicted the taxonomic origin of each gene within our identified phages. Most Absconditabacteria phages contained genes with taxonomic affiliations matching their host (Supp. Figure 6, Supp. Table 13), including one Absconditabacteria phage compatible with code 11. Five of the 57 Gracilibacteria phages contained genes predicted to originate from Gracilibacteria, including one Gracilibacteria phage that was compatible with code 11 (Supp. Figure 6, Supp. Table 13). These two analyses, in tandem with the spacer-phage matching, strongly suggest that there are, indeed, some Gracilibacteria phages and Absconditabacteria phages that are compatible with code 11.

As many of these code 11-compatible phages did not cluster coherently within the protein-sharing networks (Figure 2B, Supp. Figure 5, Supp. Figure 7) and did not contain any genes predicted to originate from their predicted host bacteria, we considered the possibility that some of the Gracilibacteria and Absconditabacteria spacer-to-phage hits might be errant, stochastic matches to the large phage databases. Such random matches would be most probable if the spacer length is unusually short. Thus, to constrain this probability, we examined the median spacer length that matched with code 11-compatible Gracilibacteria phages and Absconditabacteria phages. We found that these spacers, at 26 bp, were only slightly smaller than those that matched Saccharibacteria phages (30 bp) and those that matched clearly alternatively coded Absconditabacteria phages (28 bp), for which predicted phages generally clustered as expected (Supp. Tables 5,6). Additionally, when compared to spacers extracted from across the domain Bacteria, a spacer length of 26 bp is within the range of a typical spacer length^29^ (Supp. Figure 8).

### Phages that infect SGA and Non-SGA bacteria

We explored the host range of our putative SGA-infecting phages by comparing them to a large spacer database from a wide diversity of bacterial genomes (*Materials and Methods*). As members of Actinobacteria are known hosts of Saccharibacteria, we augmented this database with spacers from diverse Actinobacteria genomes (*Materials and Methods*). In comparing these spacers to our predicted SGA phages, we identified 23 SGA phages targeted by spacers from non-SGA bacteria (Supp. Tables 14, 15). These 23 co-targeted phages (i.e., phages targeted by both SGA and non-SGA bacteria) included 9 that potentially infect Saccharibacteria, 5 that potentially infect Absconditabacteria, and 9 that potentially infect Gracilibacteria. Seven of the nine putative Saccharibacteria phages were co-targeted by bacteria from the phylum Actinobacteria, including *Corynebacterium sp. NML130628, A. oris, Actinomyces. sp*. *HMSC075C01, A. naeslundii*, and *A. viscosus*. These species are particularly notable as a majority of cultured Saccharibacteria attach to host bacteria from the *Actinomyces* genus^5,42^. One of the co-targeted Saccharibacteria phages contained numerous spacer hits from a variety of Actinobacteria species (Figure 5A). When placed in the context of the protein-sharing network, this co-targeted phage and many other co-targeted Saccharibacteria phages are situated within a dense cluster of Saccharibacteria infecting phages (Figure 5A).

**Figure 5.**
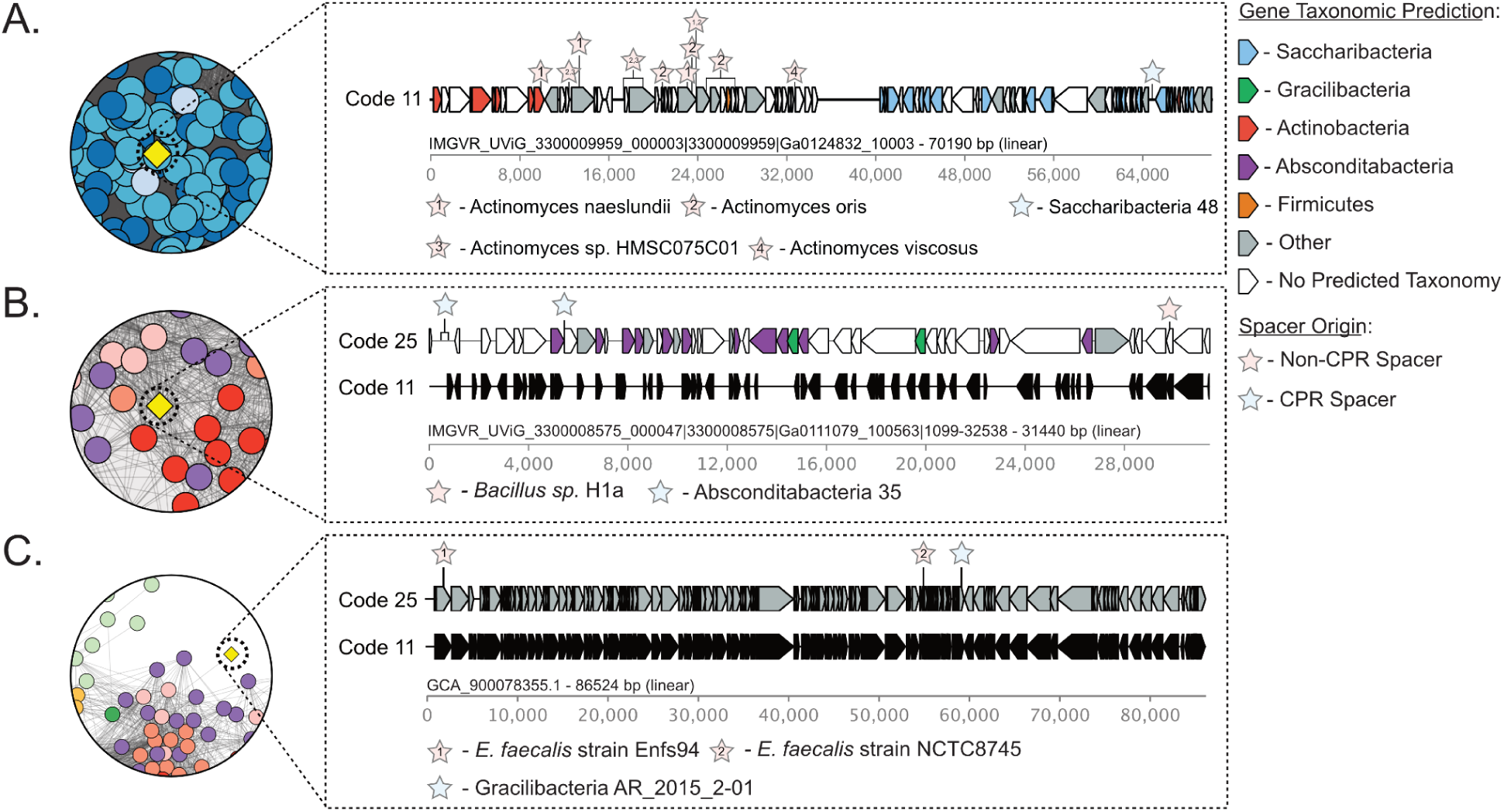
Representative Phages Targeted by Both SGA and Non-SGA Bacteria. The leftmost circular windows display a cutout of the protein-sharing network in Figure 2, with colors listed in the Figure 2 legend and the co-targeted phage spotlighted in yellow. The rightmost panels display a genome diagram of the highlighted co-targeted phage. Overlaid are gene taxonomic predictions and location of spacer matches. **A**. Saccharibacteria phage co-targeted by several *Actinomyces* species. **B**. Absconditabacteria phage co-targeted by *Bacillus sp*. H1a. **C**. Gracilibacteria phage co-targeted by *E. faecalis*.

Similarly, we also examined putative Absconditabacteria phages and Gracilibacteria phages that matched spacers from non-SGA bacteria. Three of the five putative Absconditabacteria phages matched spacers from arrays in the genomes of Firmicutes. One such phage is situated in the primary Absconditabacteria cluster within the protein-sharing network (Figure 5B). Among the nine putative Gracilibacteria phages, five have matches to spacers from arrays within Bacteroidetes species and two matched spacers from arrays within Actinobacteria species. Notably, one Gracilibacteria phage was targeted by multiple spacers from *E. faecalis* strains and was linked to the Absconditabacteria phage cluster in the protein-sharing network (Figure 5C).

By examining the genetic code of the nine Gracilibacteria phages that matched spacers from non-SGA bacteria, all nine had similar genome architectures and gene annotations in both code 11 and code 25 (representative example in Figure 5C; Supp. Tables 10, 11). Thus, they are likely capable of producing viable gene products in both their alternatively coded Gracilibacteria host and their standard coded non-SGA host. Two of the five Absconditabacteria phages that matched spacers from non-SGA bacteria, however, contained ORFs dense with in-frame TGA codons and clearly use code 25 (representative example in Figure 5B).

## DISCUSSION

Here, we examined the genetic code of SGA phages and observed that some share the genetic code of their hosts. This analysis required us to link phages to host bacteria, which we did primarily via CRISPR-Cas spacer targeting. This has been done many times previously^26–29^ and is believed to be generally robust given that the spacers in a CRISPR locus of a host bacterial genome derive directly from phage genomes^28,43^. Supporting these CRISPR spacer-based links are several lines of evidence. We observed examples of high sequence similarity between phage genes and those of their predicted host bacteria. Phages are well known to acquire genes from their hosts so the most likely explanation is that these phage genes derived from the genome of their host bacterium^44–46^. Further, in multiple protein-sharing networks, we observed strong clustering between bacteria and many of their predicted phages identified by spacer-matching. Finally, given that only a tiny subset of microbial community members use an alternative genetic code^17,19^, our linkage of alternatively coded phages to alternatively coded host bacterial groups using spacer-phage matching, as has been shown previously^20,23^, suggests that spacer-phage matches are very likely not coincidental. Thus, a variety of methods reinforce our confidence in the spacer-matching approach to identify hosts of phages.

In addition to identifying alternatively coded phages of alternatively coded Gracilibacteria and Absconditabacteria, we were surprised to identify Gracilibacteria phages and Absconditabacteria phages that lack the in-frame TGA codon usage of their hosts. These phages appear compatible with the standard code 11. The phenomenon wherein phages utilize a genetic code that is different than that of their host bacteria is not without precedent^20,21,23^. For example, some Lak phages, despite infecting standard coded bacteria of the genus *Prevotella*, have alternatively coded genomes in which the canonical stop codon TAG is reassigned to glutamine^21^. Alternatively coded phages that infect standard coded hosts may use an alternative code in part to prevent premature production of proteins that are important in late-stage infection (e.g. in-frame TGA codons within ORFs annotated as structural or lysis related proteins)^23^. To enable the translation code shift needed to produce these proteins, phage genomes often encode a suppressor tRNA that recognizes a canonical stop codon as a sense codon and incorporates a specific amino acid^20–23,47,48^. Some alternatively coded phage genomes also encode tRNA synthetases that can charge suppressor tRNAs with amino acids^23^ and release factors that terminate translation at only two of the three canonical stop codons^20,21,23^. For example, some alternative code 4 phage genomes, in which UGA is interpreted as tryptophan, encode Release Factor 1 (RF1), which only recognizes UAA and UAG, a suppressor tRNA that decodes the UGA stop codon as Tryptophan, and a Tryptophanyl tRNA-synthetase which charges the suppressor tRNA with tryptophan^23^. In combination, these previous observations underline the conclusion that phage genomes that use fewer stop codons than their host genomes require specific adaptations in the form of code shift machinery.

The situation with code 11-compatible phages, such as the Gracilibacteria phages and Absconditabacteria phages we identify in this study, is different because their lack of in-frame canonical stop codons presents no issue for translation. Where TGA is used as a stop codon, it is followed by alternative stop codons in close proximity to terminate translation. In general, this backup stop codon strategy is not uncommon in bacterial genomes and likely evolved to reduce the impact of accidental stop codon read-through^49,50^. Thus, phages that employ three stop codons should generally produce viable gene products even if the bacterial translation system only recognizes two stop codons. Unlike alternatively coded phages that infect standard coded host bacteria, phages that use the standard genetic code generally do not need to alter the translation environment of their hosts.

An intriguing finding of this study is that all identified integrated phage sequences (prophages) in Gracilibacteria and Absconditabacteria genomes were clearly alternatively coded (contained ORFs dense with in-frame TGA codons). This observation suggests that there is an advantage for a prophage to share the alternative genetic code of its host. This contrasts with the finding that some prophages adopt an alternative genetic code yet reside in bacterial genomes that use the standard bacterial code^23^. One potential explanation may be that, akin to codon optimization, higher levels of the alternative code tRNAs are expressed within alternatively coded host bacteria compared to canonical tRNAs, allowing phages with dense in-frame TGAs a more efficient translation of their gene products. The codon optimization hypothesis is supported by the high usage of TGA as a glycine codon within code 25 host bacteria^17^, that use of rare codons can lead to various translation errors^51^, and that competition over rare tRNAs can incur lower expression of gene products^52,53^.

There are two possible explanations for why the code 11-compatible Gracilibacteria phages and Absconditabacteria phages do not incorporate in-frame TGA codons. First, these standard coded phages have recently evolved to infect alternatively coded hosts. However, if this were true, we would expect such phages to be rare. As standard code compatibility is apparently not uncommon among Gracilibacteria and Absconditabacteria phages (Figures 3,4), we infer that there is an advantage for these phages to retain their standard code. Second, use of the standard code may broaden their host range, a possibility that is supported by our finding that standard code compatible Gracilibacteria phages and Absconditabacteria phages are predicted to infect standard code bacteria.

Some Saccharibacteria phages are predicted to be capable of also replicating in Actinobacteria. For cases where Actinobacteria are hosts for episymbiotic Saccharibacteria, these phages may infect both partners. Co-infecting phages thus may serve as a decoy to protect their larger bacterial symbionts from phage infection, as has been suggested previously ^31,54^.

The SGA phages reported here expand known phage diversity. Our results suggest that some of them can infect both standard and alternatively coded host bacteria, and we deduce that there is no fundamental barrier to this phenomenon. Given interest in the use of phages as therapeutics, this finding raises the possibility of producing phages to infect SGA bacteria in standard code bacteria, which may be substantially easier to cultivate than SGA bacteria themselves. Further, this may provide a path by which SGA phages can be generated for morphological characterization.

## ACKNOWLEDGMENTS

We thank Rohan Sachdeva for bioinformatic support, Adair Borges for thoughtful insights regarding alternatively coded phages, and Luis Valentin-Alvarado for support in generating genomes from the Alum Rock field site. We thank Christopher Brown for providing us access to the CRISPRbank spacer sequences and Kristopher Kieft for help with VIBRANT. This research was funded in part by the Rausser College of Natural Resources Sponsored Projects for Undergraduate Research and the Regents’ and Chancellor’s Research Fellowship (J.L). Funding was also provided by grants from the National Institutes of Health and the Moore Foundation to JFB.

## METHODS

### Absconditabacteria, Saccharibacteria, and Gracilibacteria Database Preparation

We began with a database of 864 CPR genomes derived from a previous publication^30^ that contained bacteria from three different lineages: Absconditabacteria, Saccharibacteria, and Gracilibacteria. We de-replicated the database using dRep^55^ at 99% ANI clustering and default alignment fraction (10%). For each genome, we predicted protein sequences using the “single” mode of Prodigal^56^. For Saccharibacteria genomes, genes were predicted in genetic code 11. As Gracilibacteria and Absconditabacteria adhere to a non-standard genetic code, code 25^17,18^, Gracilibacteria and Absconditabacteria genes were predicted in genetic code 25. Genome annotation was performed using hmmsearch from HMMER3^57^ with the KEGG^58^, UniRef100^59^, and UniProt^60^ databases. Gene taxonomic predictions were performed using USEARCH^61^ with the UniRef100 database.

A phylogenetic tree of the nonredundant genomes was constructed, as previously described^30^, using a concatenated set of 16 syntenic ribosomal proteins. Briefly, sequences were individually aligned using MAFFT^62^, trimmed using BMGE^63^, and concatenated. A maximum-likelihood tree was then inferred for the concatenated alignments using IQ-tree^64^ (ultrafast bootstrap, -bb 1000, -m MFP) and visualized with iTOL^65^.

### CRISPR-Cas Array Prediction and Curation

To search for CRISPR arrays in the SGA genome database, we ran CRISPRCas Finder^66^ (CCF) on all genomes. We then selected scaffolds containing CRISPR arrays designated as evidence level 3 or 4 - arrays deemed highly likely candidates by CCF - for further curation. We manually curated the scaffolds containing high evidence level CRISPR arrays to ensure they did not originate from misbinning. Our manual curation considered three complementary metrics: we considered a scaffold to be from SGA bacteria if 1) the majority of predicted proteins appeared to have the closest taxonomic hits to SGA bacteria, 2) if individual, phylogenetically informative proteins appeared to have the closet taxonomic hits to SGA bacteria, and 3) if scaffolds displayed high coding density in code 25 relative to code 11.

### Identification of Complete CRISPR-Cas Systems

To identify complete CRISPR-Cas systems present in our database, among the genomes containing high-confidence, manually curated CRISPR arrays, we searched for Cas proteins using the full suite of TIGRfam HMMs^67^ (hmmsearch, model-specific noise cutoff). We additionally manually curated all scaffolds containing Cas gene annotations to ensure they were from SGA bacteria using the metrics described above. We defined complete CRISPR-Cas systems based on previously published descriptions of various CRISPR-Cas systems^32,36^: For each array, 1) if *cas9, csn2, cas1*, and *cas2* genes were also encoded within in the same genome, we categorized it as a type II-A system; 2) if *cas9, cas1, and cas2* genes and no *csn2* genes were encoded within the same genome, we categorized it as a type II-C1 system; 3) if *cpf1, cas1, cas4*, and *cas2* genes were encoded in the same genome, we categorized it as a type V-A system; 4) if *cas10, cas7, cas5*, and *csm2* genes were encoded in the same genome, we considered it a complete type III-A system; and 5) if *cas10, cas7, cas5*, and *cmr5* genes were encoded in the same genome, we considered it a complete type III-B system. Complete CRISPR-Cas systems were visualized using gggenes (https://github.com/wilkox/gggenes). For CRISPR-Cas systems containing all Cas proteins on the same scaffold, AAI similarities and Cas operon architectures were visualized using Clinker^68^ at default parameters.

To assess the novelty of identified Cas genes, we compared the Cas genes within each complete CRISPR-Cas system to the NCBI nr database using BLASTp (evalue ≥ 1e-3, coverage ≥ 0.75) and retained the best hit per gene based on percent identity.

### Compiling a Saccharibacteria, Absconditabacteria, and Gracilibacteria Spacer Database

To compile a spacer database, we extracted all spacers from high evidence level arrays on scaffolds from our redundant SGA database (862 genomes) that passed our manual curation step. In addition, we ran a previously described spacer array expansion step to gather additional spacers from variant sequences that are not reflected in the consensus metagenomic assembly^38^. Briefly, if available, we gathered the metagenomic reads originally used to assemble each genome. We then reassembled the reads using MEGAHIT at default parameters^69^, mapped reads back to assembled contigs using Bowtie2 at default parameters^70^, and predicted proteins using the ‘meta’ flag of Prodigal. We compared the newly assembled scaffolds to the original, publicly available scaffolds. If a newly assembled scaffold matched an original, manually-curated scaffold above the thresholds of 95% coverage and 90% ANI, we predicted CRISPR arrays in the newly assembled scaffold using CCF, extracted spacers from high-evidence level arrays, and added extracted spacers to our spacer database. We de-replicated the spacer database at 100% ANI using USEARCH.

### Identification of CPR-infecting Phages

To search for phages putatively infecting SGA bacteria, we compared each unique spacer in our spacer database to two phage databases: IMG/VRv3^26^ and GVD Human Gut Virome^39^. BLASTn parameters were set to at least 95% coverage of the spacer and 1 or less allowed mismatch with the specific flags: -task “blastn-short” -word_size 7 -gapopen 10 -gapextend 2 -penalty -1. Prophages within the CPR genomes were predicted using VIBRANT at default parameters^71^.

### Absconditabacteria, Saccharibacteria, and Gracilibacteria Phage Characterization

To dereplicate the putative SGA-infecting phages, we ran dRep at 99% ANI clustering and default alignment fraction (10%). We additionally predicted phage genome circularization using VIBRANT.

To identify the likely genetic code of putative SGA-infecting phages larger than 20 kb, we used the Prodigal ‘single’ flag to calculate the coding density of each phage in genetic codes 11 and 25. Further, genome diagrams of Gracilibacteria and Absconditabacteria prophages and phages greater than 20 kb were generated using the Prodigal ORF predictions in code 11 and code 25. In-frame TGA codons were additionally located within the ORF predictions. The genome diagrams of the phages in both code 11 and code 25 were visualized using gggenes.

To annotate phage proteins, we used the Prodigal gene predictions in the genetic code 11 for putative Saccharibacteria phages, and the Prodigal gene predictions in genetic code 25 for putative Gracilibacteria phages and Absconditabacteria phages. We annotated predicted proteins using Kegg, UniProt, and pVOG^72^ HMM profiles with hmmsearch from HMMER3. Gene taxonomic predictions were performed using DIAMOND^73^ with the UniRef100 database.

To compare the putative SGA-infecting phages to reference phages and their predicted host bacteria, we constructed two protein-sharing networks using vContact2^74^ (--rel-mode Diamond, –vcs-mode ClusterONE, --pcs-mode MCL). One network linked the proteomes of the putative SGA phages (including prophages) identified in this study, previously identified CPR phages^23,26,40,41^, and SGA bacteria. The second network linked the SGA phages identified in this study, the previously identified SGA phages, and non-SGA reference phages (--db ‘ProkaryoticViralRefSeq201-Merged’). The resulting protein-sharing networks and their associated metadata were visualized in Cytoscape^75^.

### Host Range of CPR-infecting Phages

To examine the host range of putative CPR-infecting phages, we compared spacers from four comprehensive databases^41,66,76^ composed of spacers from across the domain Bacteria to the predicted SGA-infecting phages. As before, the BLASTn parameters were set to at least 95% coverage of the spacer and 1 or less allowed mismatch with the specific flags: -task “blastn-short” -word_size 7 -gapopen 10 -gapextend 2 -penalty -1.

We additionally constructed a database for Actinobacteria, some of which are known hosts of Saccharibacteria, by sampling one genome per species-level group from GTDB (release 95, August 2020). Using a similar workflow as with the SGA database, we searched these genomes for high evidence level arrays with CCF, extracted spacers, and compared them to the putative CPR-infecting phages with the above parameters. Visualization of spacer hits was performed using DNA Features Viewer^77^.

## DATA AVAILABILITY

Non-redundant SGA genome accessions and associated metadata are listed in Supp. Table 1. Redundant SGA genomes will be made available on Zenodo. SGA phage accessions are listed in Supp. Tables 4 and 5, including IMG/VR UViGs and GVD scaffold names. All code used in this project will be made available on Github.

## SUPPLEMENTAL FIGURES

**Supp. Figure 1.**
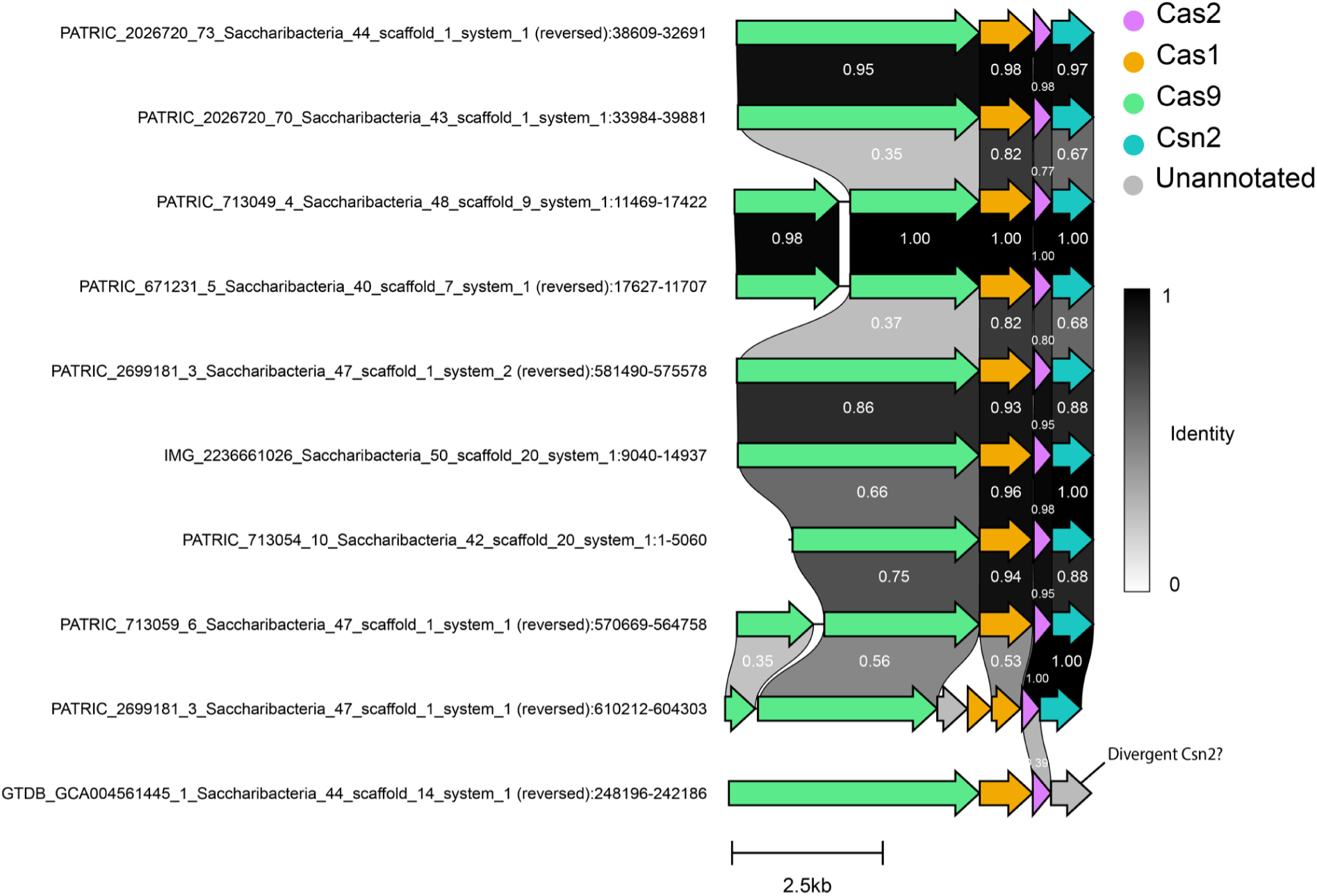
Shared Amino Acid Identity and Architecture of Identified Type II-A CRISPR-Cas Systems. Scaffolds containing a complete Type II-A Cas locus were compared by AAI. Scaffolds were additionally manually curated to avoid contamination. Order was determined based on cluster similarity. Only the best AAI links between ORFs are shown.

**Supp. Figure 2.**
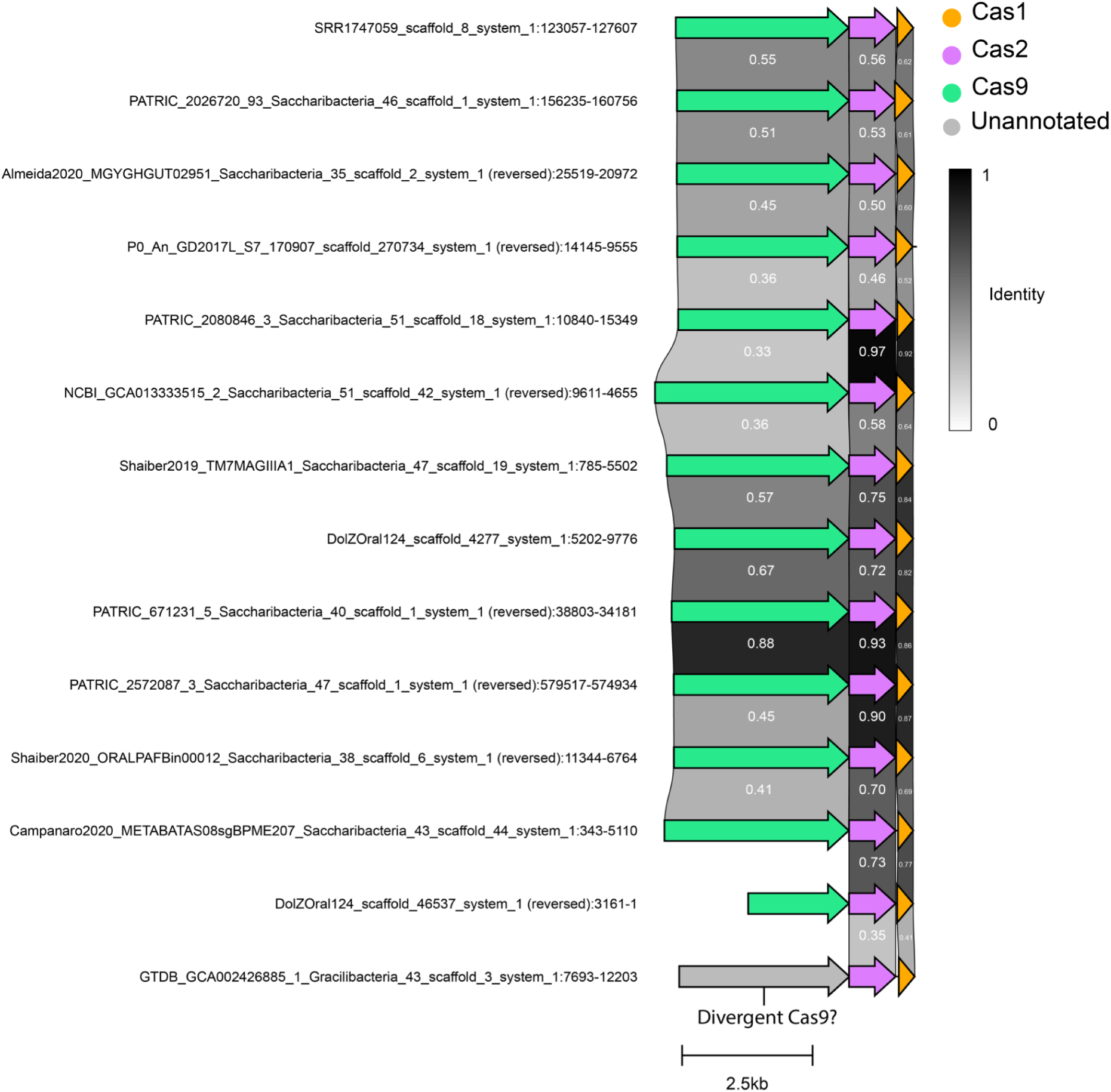
Shared Amino Acid Identity and Architecture Variation of Identified Type II-C CRISPR-Cas Systems. Scaffolds containing a complete Type II-C Cas locus were compared by AAI. Scaffolds were additionally manually curated to avoid contamination. Order was determined based on cluster similarity. Only the best AAI links between scaffolds are shown.

**Supp. Figure 3.**
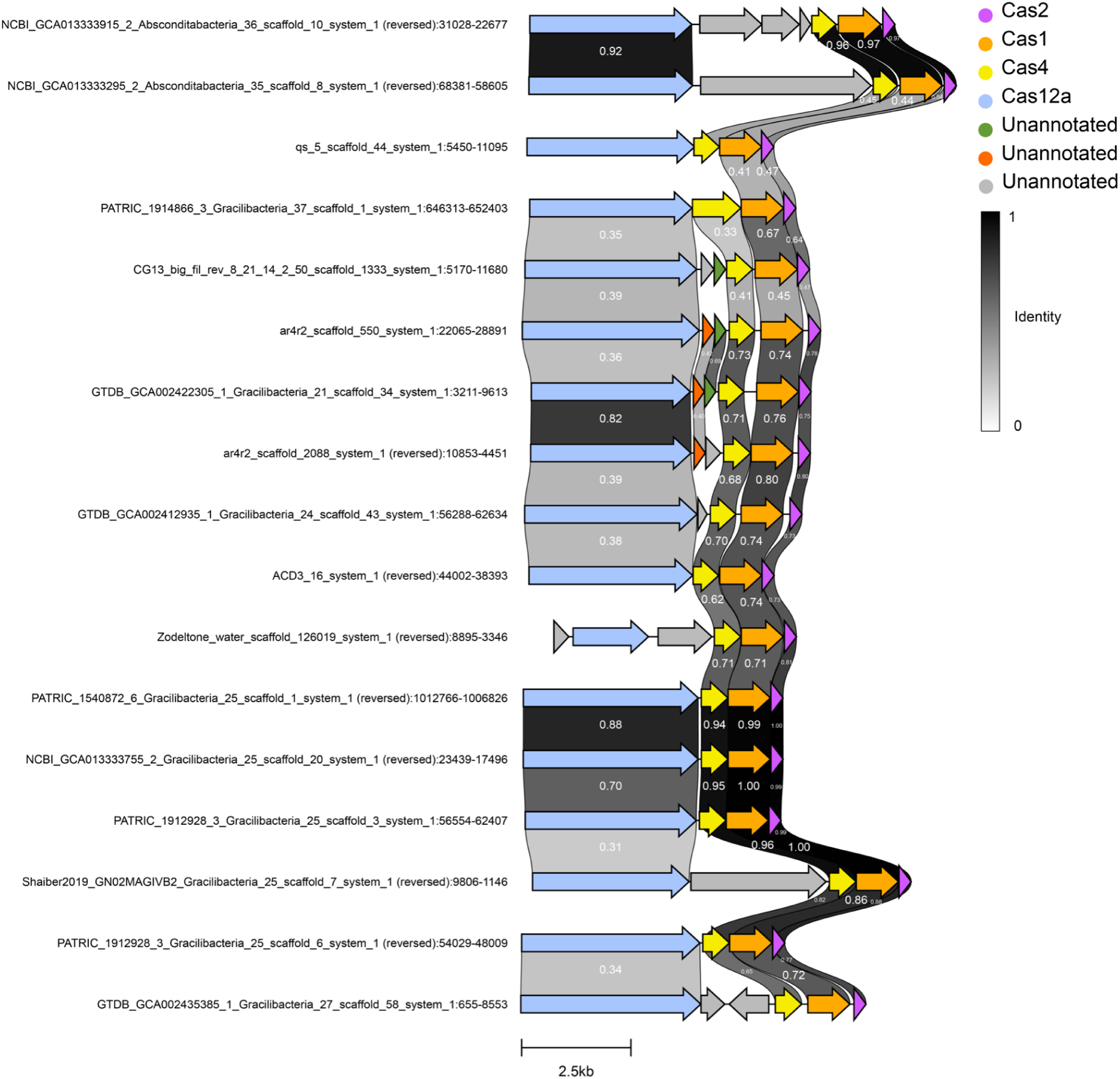
Shared Amino Acid Identity and Architecture Variation of Identified Type V-A CRISPR-Cas Systems. Scaffolds containing a complete Type V-A Cas locus were compared by AAI. Scaffolds were additionally manually curated to avoid contamination. Order was determined based on cluster similarity. Only the best AAI links between scaffolds are shown.

**Supp. Figure 4.**
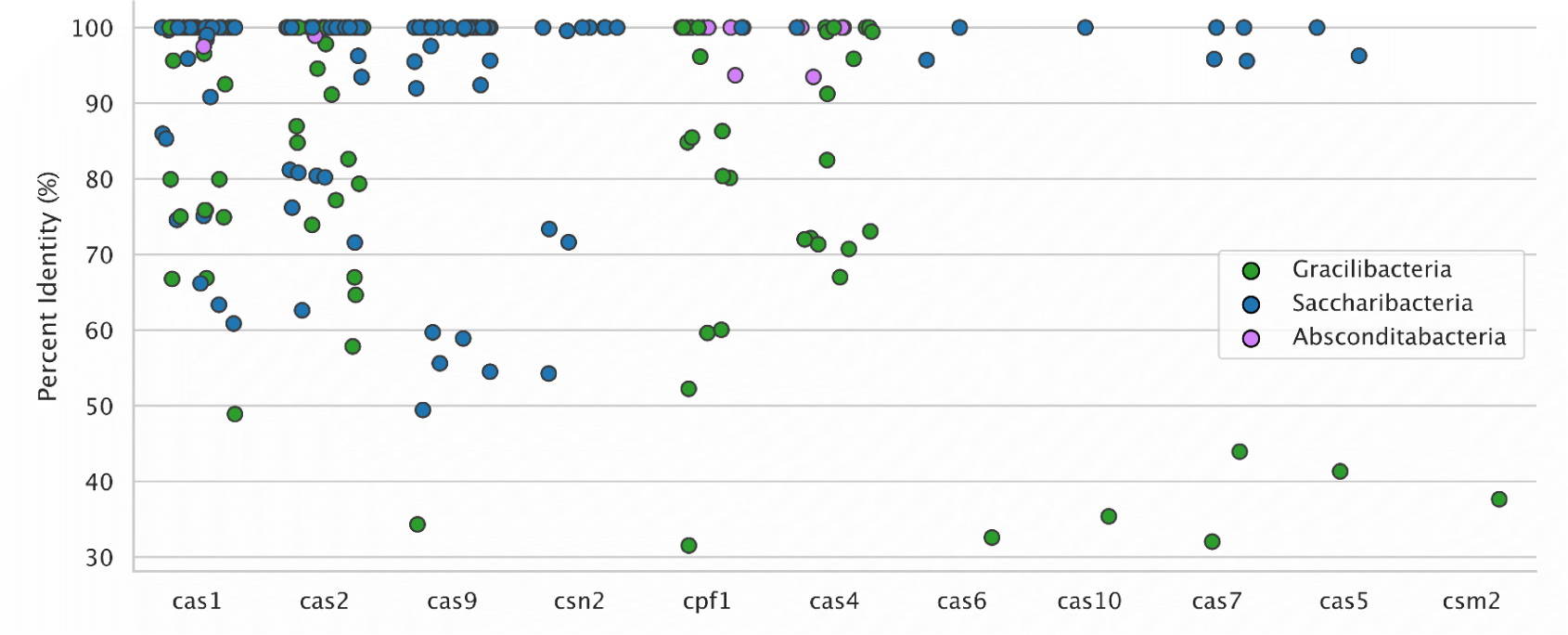
BLASTp of Cas genes from our SGA genome database to the NCBI nr database. Each dot represents a Cas gene from our non-redundant SGA genome database positioned based on their highest percent identity hit to the NCBI nr database. A minimum threshold of 75% coverage of a reference sequence was employed. Node colors denote the SGA lineage of the genome encoded with the gene.

**Supp. Figure 5.**
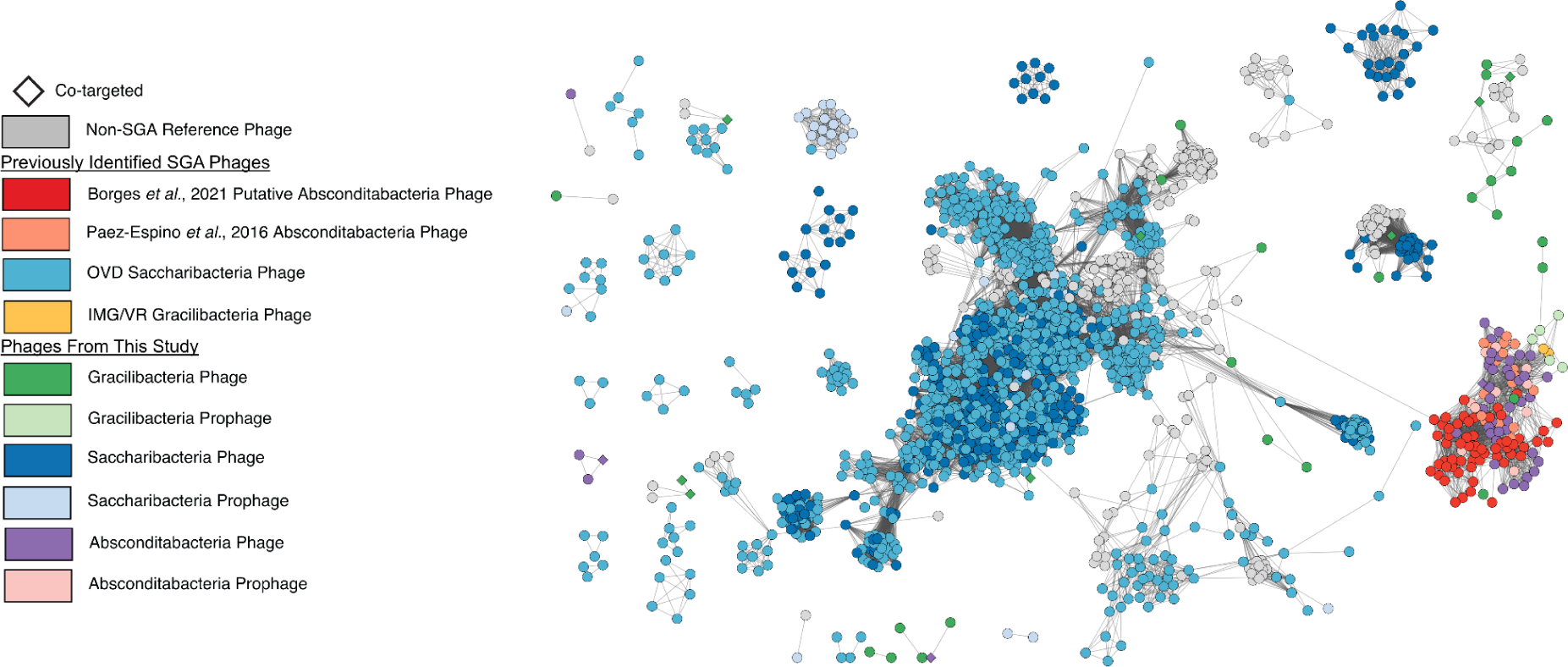
Protein-sharing network of the putative SGA phages, where each node represents a phage and nodes are clustered together based on protein similarity and number of shared proteins. Non-CPR reference phages and previously identified SGA phages were included^23,40,41^ in the network. Only reference phages with at least one neighboring SGA phage are displayed. Phage nodes are colored based on the predicted host of the phages. Co-targeted phages indicate those targeted by spacers from CRISPR-Cas arrays of both SGA and non-SGA bacteria.

**Supp. Figure 6.**
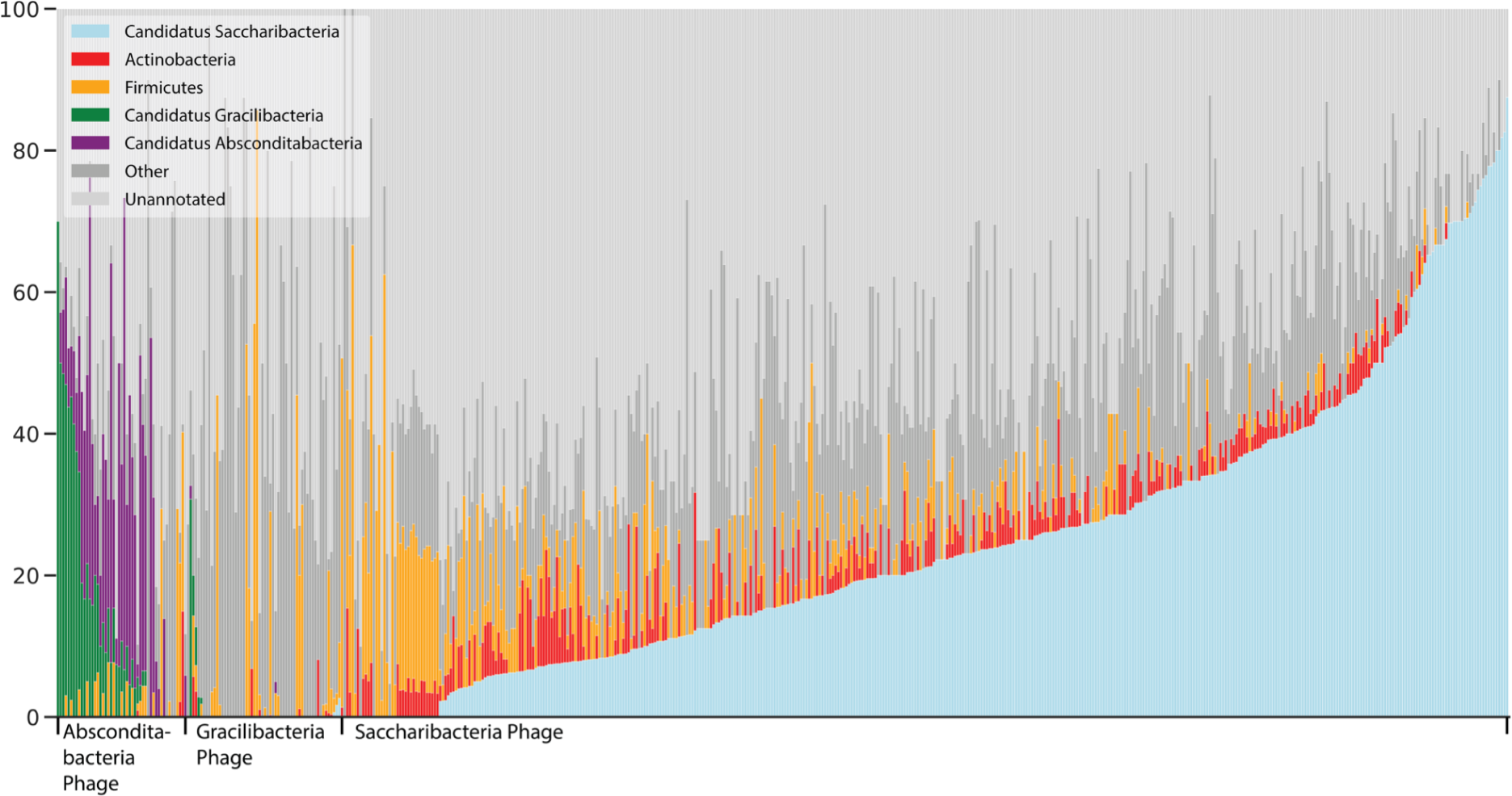
DIAMOND taxonomy predictions for putative SGA phage genes. The thresholds “e-value < 10^−6^ and coverage ≥ 70%” were used to filter results. Only the best hit to each gene based on bitscore was retained.

**Supp. Figure 7.**
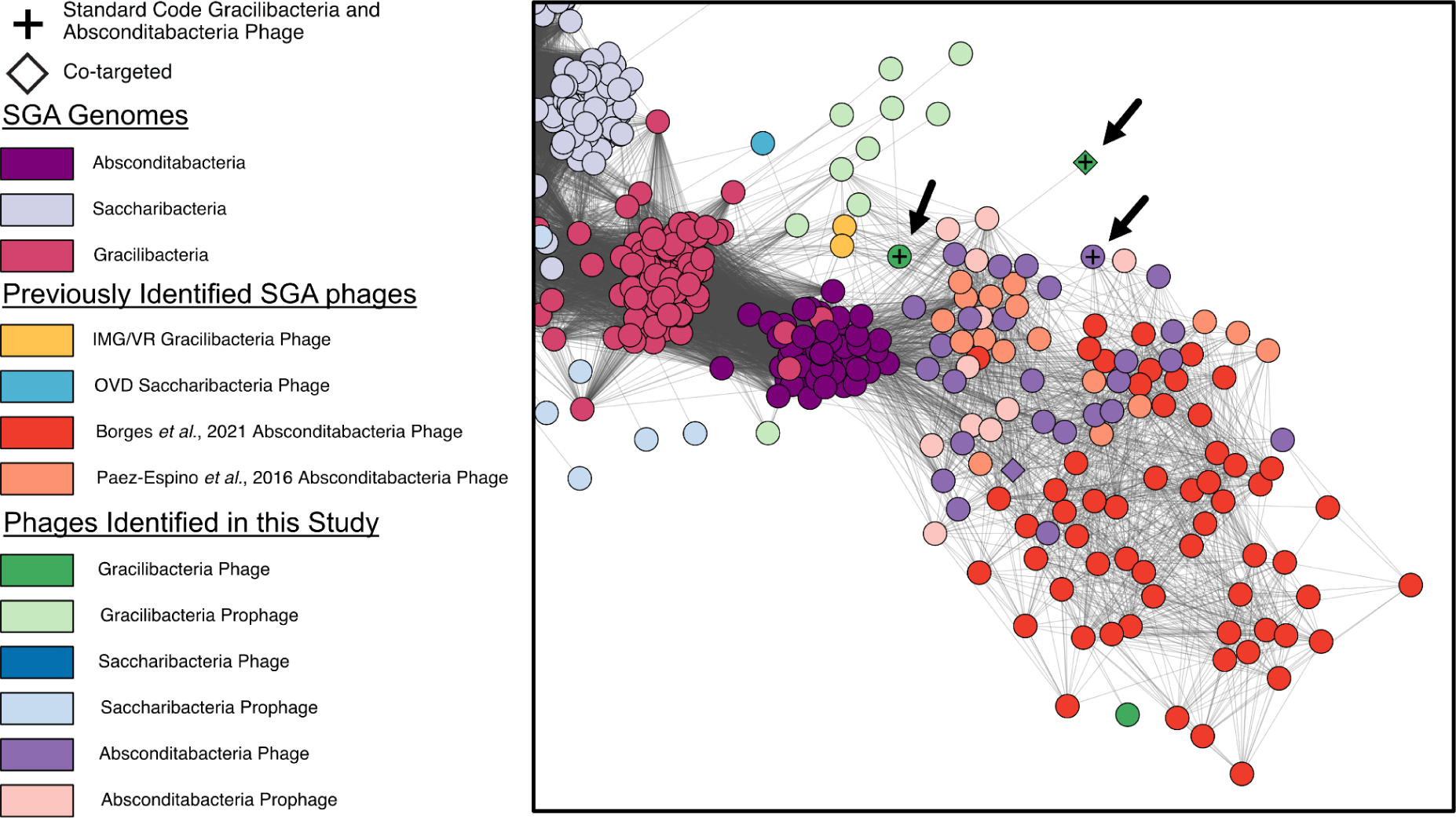
Close up of the protein-sharing network presented in Figure 2. Arrows indicate predicted Gracilibacteria phages and Absconditabacteria phages that are compatible with genetic code 11.

**Supp. Figure 8.**
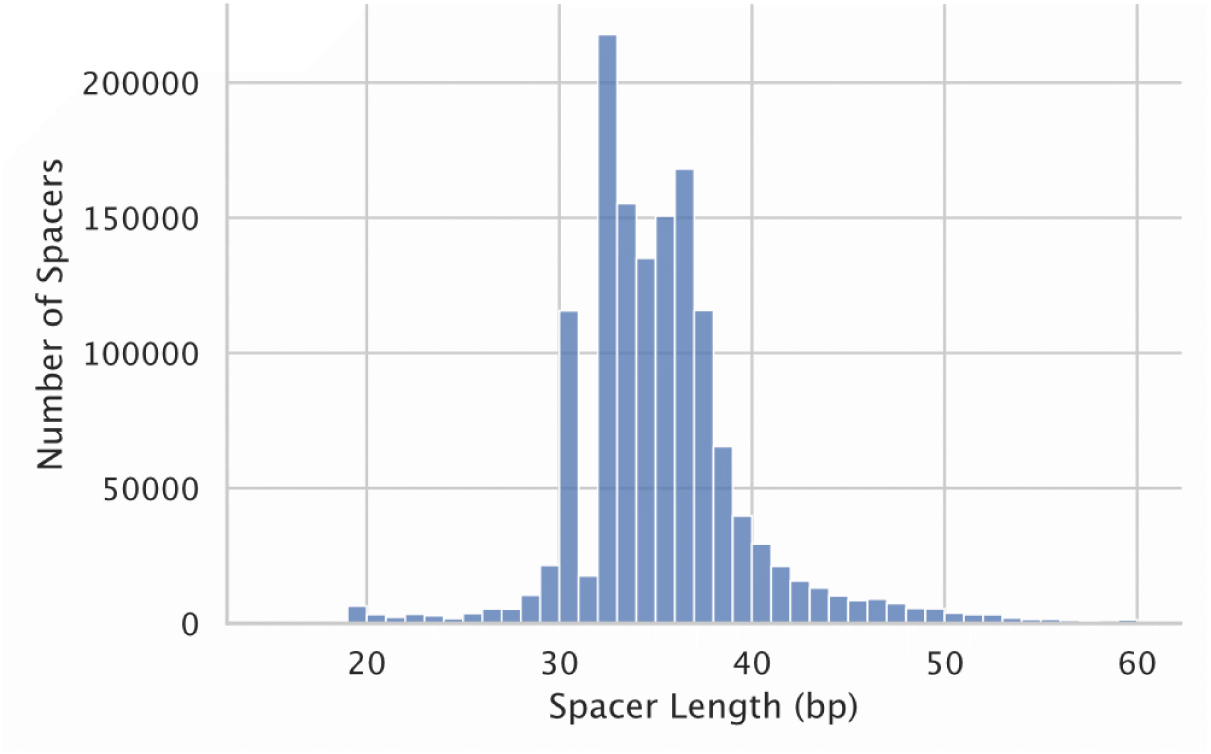
Histogram of iPHoP spacer length. All spacers from iPHoP^29^ were extracted and binned based on their length.

